# Many paths to the same goal: metaheuristic operation of brains during natural behavior

**DOI:** 10.1101/697607

**Authors:** Brian J. Jackson, Gusti Lulu Fatima, Sujean Oh, David H. Gire

**Affiliations:** Department of Psychology, University of Washington, Seattle, WA 98195, USA

## Abstract

During self-guided behaviors animals rapidly identify the constraints of the problems they face and adaptively employ appropriate cognitive strategies and heuristics to solve these problems^1,2^. This ability is currently an area of active investigation in artificial intelligence^3^. Recent work in computer science has suggested that this type of flexible problem solving could be achievable with metaheuristic approaches in which specific algorithms are selected based upon the identified demands of the problem to be solved^4,5,6,7^. Investigating how animals employ such metaheuristics while solving self-guided natural problems is a fertile area for biologically inspired algorithm development. Here we show that animals adaptively shift cognitive resources between sensory and memory systems during natural behavior to optimize performance under uncertainty. We demonstrate this using a new, laboratory-based discovery method to define the strategies used to solve a difficult optimization scenario, the stochastic “traveling salesman” problem^5,8,9^. Using this system we precisely manipulated the strength of prior information available to animals as well as the complexity of the problem. We find that rats are capable of efficiently solving this problem, even under conditions in which prior information is unreliable and the space of possible solutions is large. We compared animal performance to a Bayesian search and found that performance is consistent with a metaheuristic approach that adaptively allocates cognitive resources between sensory processing and memory, enhancing sensory acuity and reducing memory load under conditions in which prior information is unreliable. Our findings set the foundation for new approaches to understand the neural substrates of natural behavior as well as the rational development of biologically inspired metaheuristic approaches for complex real-world optimization.

## Main Text

The ability to rapidly solve complex problems is a defining feature of animal intelligence. Indeed, varieties of animals solve difficult optimization problems nearly instantaneously^10,11,12,13^. Metaheuristics, where an agent selects a heuristic to solve a difficult optimization problem especially when information is imperfect, are a promising computational framework that could support such complex problem solving^14,15^. However, it is not known whether animals use this type of reasoning when solving challenging optimization problems.

Here we investigated how rats solve a notoriously difficult optimization scenario, the stochastic traveling salesman problem. In this problem, an agent must establish the most efficient (i.e. shortest) route between targets with only probabilistic information regarding whether targets will be present or absent at each location^16,17^. We adapted this problem for experimental investigation through the use of an automated system for precise, computer-controlled food pellet placement within a large foraging arena (Fig. 1a). We divided a cohort of 12 rats into 3 equal groups that foraged within environments of high, medium, and low food location predictability (Fig. 1b). Animals in each group were tested across precisely replicated pellet placements (Fig. 1c) and all placements used had equivalent optimal path lengths, as calculated through a genetic algorithm solution to the traveling salesman problem for each pellet placement (Fig. 1d, and see methods). We generated sequences of pellet locations over days to create distributions that were extremely well-predicted by prior experience as well as distributions that were unable to be anticipated based upon prior pellet locations. To generate pellet placements with controlled levels of predictability we quantified the between trial minimum distance for each pellet of a given distribution and all pellets of the previous trial’s distribution and set this value to be low for the computer-generated set of locations used for predictable conditions and to be high for the unpredictable condition (Fig. 1e). The lower values for pellets in predictable distributions indicate that these pellets are in areas that are extremely close to where pellets were located on the previous trial, allowing animals to create an expectation over repeated searches. This is also demonstrated through a reduction of the relative entropy (a measure of surprise) of newly-encountered pellet distributions following multiple days of training for animals in high and medium predictability conditions. Animals could not develop such an expectation under low levels of predictability and relative entropy does not decrease with training for the unpredictable distribution (Fig. 1f). In all conditions, animals searched for an average of 7 pellets, with the precise number on a given trial unknown to the animal (Fig. 1g). This results in typically 7!, or 5,040 possible sequences of pellet acquisition, with most sequences being extremely sub-optimal. All animals favored a small subset of near-optimal acquisition sequences (Fig. 1h), consistent with findings in non-probabilistic optimization across a number of species^18^.

**Fig. 1:**
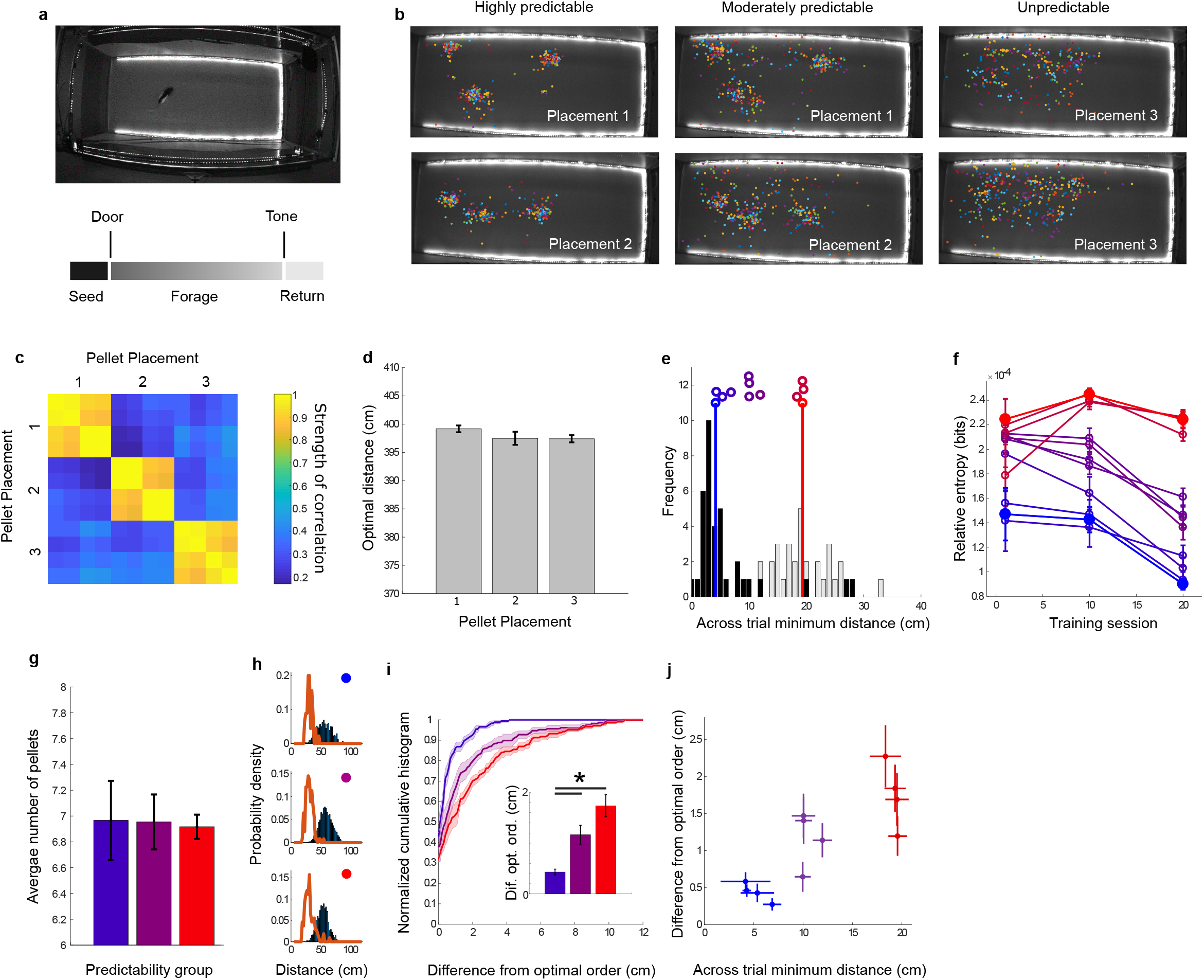
A computer-controlled probabilistic traveling salesman task enables direct tests of behavioral strategies under uncertainty. **a) (top)** A large, automated arena with a rat shown for scale. (bottom) The temporal structure of a typical trial. **b)** Rats forage for pellets in highly predictable (left), moderately predictable (center), and actively randomized (right) pellet placements. Placements are shown across all trials (20 days, 3 trials per day). **c)** The automated system allows for reproducible pellet placement across animals. From the top to bottom of the matrix correlation coefficients are shown for two different predictable distributions and the single unpredictable distribution. **d)** Pellet distributions from each placement shown in panels b) and c) have equivalent optimal path lengths. **e)** Example histograms are shown for the most predictable (black) and least predictable (gray) distributions that were tested. Vertical colored lines show the mean for the predictable (blue) and unpredictable (red) distributions. The distributions for all animals are plotted as colored circles, with color corresponding to across trial minimum distance. **f)** Relative entropy for each predictability grouping (high, blue, medium purple, and randomized red) across sessions of training. Higher values indicate higher entropy. **g)** Average number of pellets per trial for each predictability level. h) Average distance per pellet. Rats acquire pellets in a sequence that is extremely efficient (red lines) compared to a random sampling of all possible sequences (blue bars). Predictability decreases from top to bottom. **i)** Cumulative histograms of difference between animal acquisition sequence and optimal sequence on each trial. Large positive values indicate less effective pellet acquisition sequences. **j)** Scatter plot showing the relation between predictability of distribution (x axis) and difference between animal acquisition sequence and optimal sequence (y axis) for all 12 animals.

However, animals in the lowest predictability group showed a reduced ability to acquire pellets in optimal sequences (Fig. 1i), with animals trained with highly predictable distributions (blue) choosing the most efficient sequences while animals trained with unpredictable distributions (red) followed significantly less efficient acquisition sequences (highly predictable shorter path length than moderately predictable (p<0.05) and unpredictable (p<0.05), see inset of Fig. 1i; n = 4 animals per predictability group, see methods for statistical tests used for all comparisons). This is observed across the entire cohort of animals as a significant positive correlation between the unpredictability of a distribution and the amount of error relative to an optimal path (Fig. 1j; R = 0.83; p <0.001, n = 12 animals).

Nearest neighbor tours are a common strategy used to solve the traveling salesman problem^19,20^. Under this strategy, the agent simply travels to the next nearest target location until all targets have been visited. While not optimal, this approach is computationally simple, resulting in rapid solutions with time to solve scaling well with task complexity. We found that the nearest neighbor path is close to optimal across all distributions, though it occasionally results in sub-optimal paths (Fig. 2a). Consistent with a nearest neighbor strategy, animals were more likely to follow an optimal pellet acquisition sequence on trials in which a nearest neighbor tour resulted in an optimal sequence (percent of tours that are optimally ordered 18.1 +/− 1.6%) than on trials in which the nearest neighbor tour was sub-optimal (percent of tours that are optimally ordered 1.9 +/− 0.3%; Fig. 2b; p<0.0001; n = 12 animals). In our task, which involves probabilistic presence of pellets, this strategy can be implemented through two different heuristics: in a sensory-guided heuristic animals use cues (odor or vision) to navigate towards the nearest detected target; in a memory-guided heuristic animals use prior information to navigate towards the nearest, most likely locations of pellets. We next investigated which of the two alternative heuristics might guide a nearest neighbor search within each level of uncertainty. Over training, animals across all predictability levels significantly increased their probability to travel to the nearest pellet during search. However, the number of days of training taken for this to occur was dependent upon the predictability of the pellet distribution (Fig. 2c; significant improvement on days 2-5 for highly and moderately predictable locations, significant improvement not until days 10-15 for unpredictable locations; p <0.05 compared to day 1, n=4 for all groups). We found that animals searching in highly predictable distributions were the most effective at navigating to nearest neighbor locations and that the advantage of predictability was strongest for large distances (0.5-1 meter), much beyond the range of possible sensory cues^21^ (see Supp. Fig. 1). Animals foraging in uncertain environments without a high level of predictability showed a dramatic reduction in the ability to find the nearest neighbor location when searching over these large distances (Fig. 2d; 84.9 +/− 1.4% trajectories from long distance are to nearest neighbor in predictable distributions, 68.3 +/− 2.6% in unpredictable distributions, p<0.05, n = 4 animals in each group). The impact of traveling from a distance was significantly more negative for animals in unpredictable distributions (7 +/− 2% reduction from short to long distance in predictable distributions, 20 +/− 2% in unpredictable distributions, p < 0.05, n = 4 animals per group). In addition to supporting better-ordered search routes (Fig. 1j) and travel to the nearest target from farther away (Fig. 2d), predictable distributions also enabled rats to enhance the paths taken between pellets. After training, the efficiency of the paths taken between pellets was significantly correlated with the predictability of the distribution (Fig. 2e; efficiency negatively correlated with between trial distance; R = −0.79; p = 0.0022; n = 12).These results are consistent with animals employing two distinct heuristics to find the next nearest pellet. In a sensory-dominated heuristic animals approach the nearest sensed pellet, while in a memory-dominated heuristic animals approach the nearest remembered location, enabling more efficient, planned routes to emerge.

**Fig. 2:**
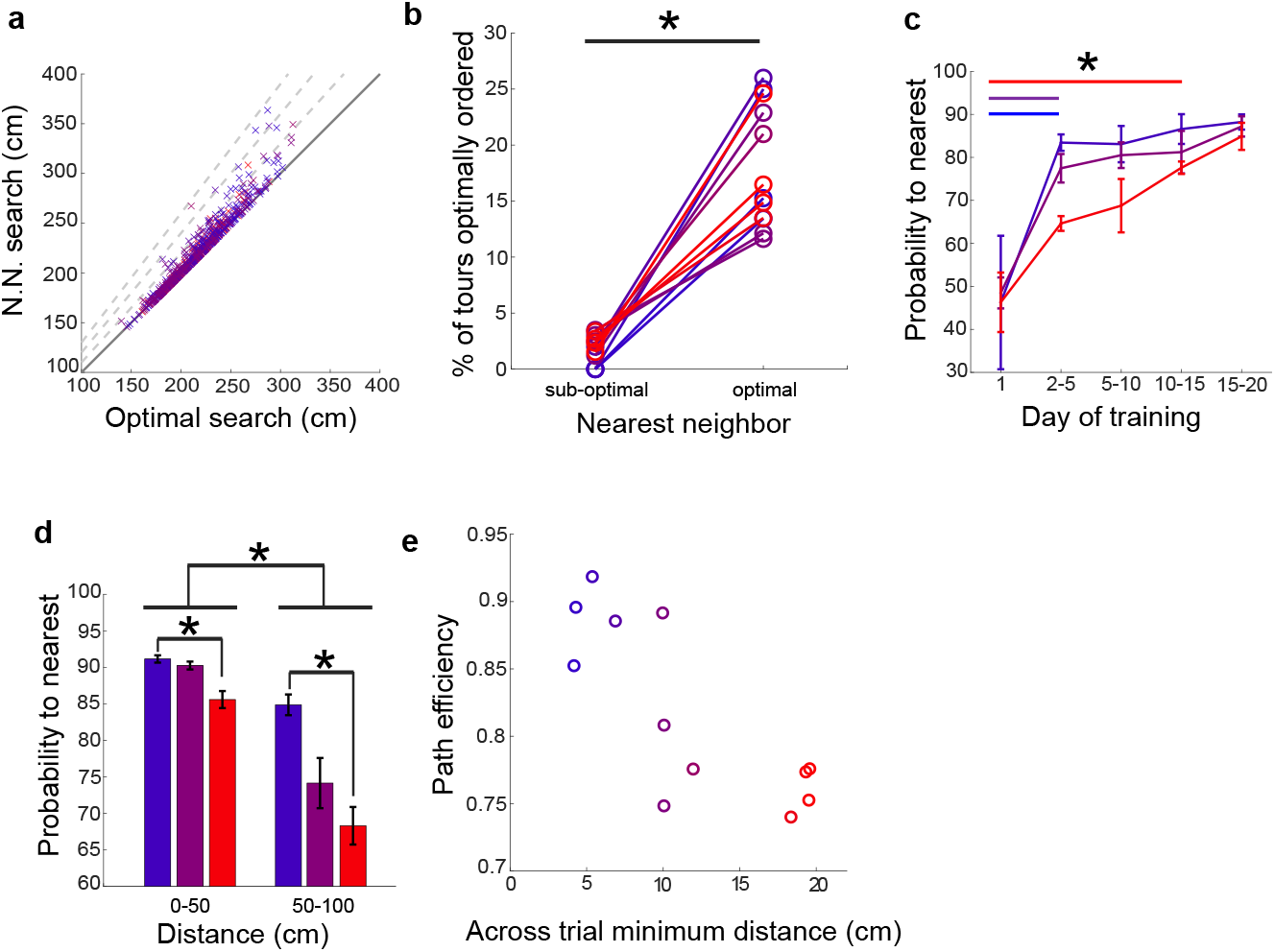
Rats develop multiple heuristics to support a nearest neighbor search. **a)** Performance of a nearest neighbor strategy on all distributions tested in this study when compared to the optimal path length. Dashed lines represent 10, 20 and 30% above optimal. **b)** Animal performance on trials in which a nearest neighbor search is optimal vs. trials in which a nearest neighbor search is sub-optimal. **c)** The probability that rats in each predictability group acquire the nearest pellet during search increases during training for all groups. **d)** Probability that an animal will acquire the next nearest pellet as a function of distance to that pellet with groupings of short (<50 cm) and long (>50 cm) distances. **e)** Efficiency of paths taken between pellets is shown as a function of the predictability of each series of pellet distributions upon which animals were trained.

We next sought to more precisely quantify the role of sensory information and memory in the navigation strategies used by animals under varying levels of uncertainty. To perform this analysis we modeled animal behavior as a Bayesian search with multiple free parameters^22,23^. These parameters include the length of memory for the prior, the distance over which sensory signals from the pellets are detected and the relative weighting of sensory and memory terms. We allowed these parameters to vary on a multidimensional grid and analyzed goodness of fit to actual animal performance as the correlation between trial-by-trial performance of the simulated searcher and the animal (see methods). As expected, searches with long-range, noiseless sensory information lead to a perfect nearest neighbor search and do not correlate well with animal behavior since rats do not have access to perfect information and need to use local sensory information or learned locations to navigate (see Supp. Fig. 1). Similarly, searches with only a memory term also do not correlate well with actual behavior (Fig. 3a; see Supp. Fig. 2 for a more extensive exploration of parameter space). Consistent with animals under different levels of uncertainty using diverse search strategies, we found that any set of a wide range of parameters applied uniformly to all animals resulted in only moderate correlation with actual behavior (Supp. Fig. 2b). We next allowed parameters to vary individually for each animal. While this approach will trivially result in a better fit due to the increased number of free parameters (Fig. 3a, p<0.01; n= 12), Supp. Fig. 2b), we used the values of parameters obtained for these individual fits to examine the contribution of sensory and memory input to the Bayesian search that best matched each animal’s performance. When varying the length of memory used by the searcher we found that Bayesian searches across the most predictable distributions benefited from increased memory with an increase in correlation to actual animal performance when the simulated searcher had access to cumulative memory of previous searches (predictable, single trial memory: R = 0.12 +/− 0.05; cumulative memory R = 0.66 +/− 0.03; p<0.05; n = 4). Searches across moderately predictable and unpredictable distributions did not show a significant increase in correlation with animal behavior with increased memory (Fig. 3b). Consistent with these results, the impact of shuffling prior distributions on agent performance was directly related to the predictability of the data set (Supplemental Figure 2c). To quantify the impact of sensory input on Bayesian searches we combined the weighting given to sensory input with the distance from which each agent could detect a target to create a measure of sensory acuity for each simulated agent (see methods). This measure was well correlated with increasing relative entropy of the training set, suggesting that animals increased sensory acuity under uncertainty (Fig. 3 c, upper panel; R = 0.8469; p = 0.005). We also used the length of memory for the best match to animal behavior to create a metric for long-term memory usage (see methods). We found a significant inverse correlation between relative entropy and long-term memory usage (Fig. 3c, lower panel; R = −0.7252; p = 0.0076), suggesting that as the training set became more predictable animals relied more on long-term memory. Our results are consistent with a metaheuristic strategy that increases sensory acuity and reduces memory load in direct relation to the level of uncertainty in an environment (Fig. 3d). This increased reliance on sensory input allows animals searching across unpredictable environments to employ an effective nearest neighbor strategy with nearly the same efficacy as animals that are operating in highly predictable environments, although a sensory-guided heuristic fails at long distances. Conversely, animals operating in predictable environments reduce their reliance on sensory input in favor of stereotyped and efficient searches based on long-term memory, which allows them to enhance search tours over long distances. Taken together, these results suggest that animals employ uncertainty-based metaheuristics to select appropriate strategies to allocate cognitive resources and solve complex natural problems.

**Fig. 3:**
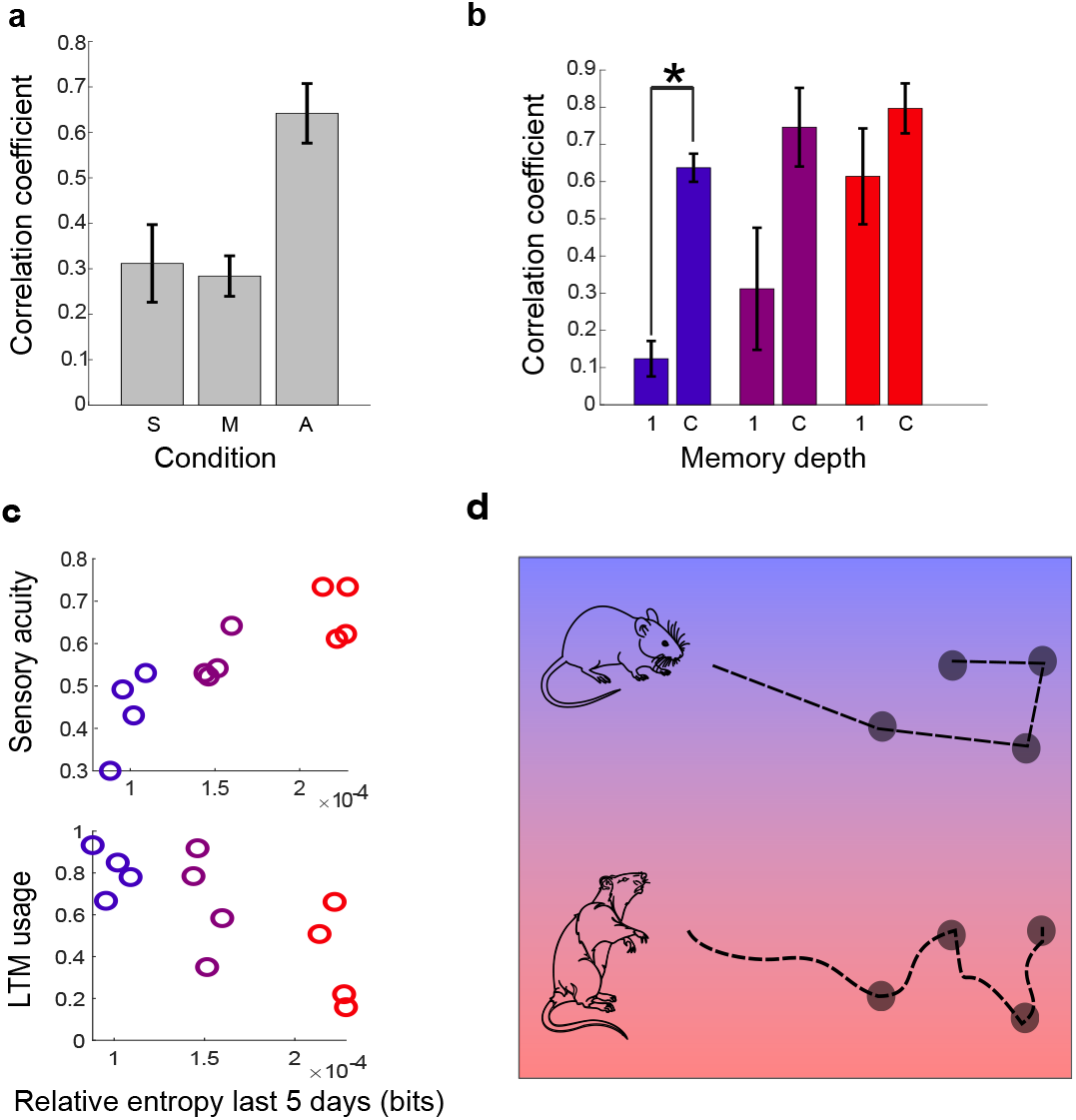
A Bayesian search with adaptive sensory acuity and memory depth explains performance under uncertainty. **a)** Correlation to animal performance of models with parameters emphasizing sensory (S) or memory (M) guidance or an adaptive model (A) individually fit to each animal. **b)** Correlation between animal behavior and a Bayesian search with either single trial memory (1) or best performance with cumulative memory (C). **c) (top)** Sensory acuity based on the best fit Bayesian search parameters vs. relative entropy based on the distributions that animals have experienced. **(bottom)** Long-term memory usage vs. relative entropy of pellet distributions encountered. **d)** A schematic of the main results, showing that animals adaptively change the heuristics used for a search depending upon the level of uncertainty of the environment, here depicted as a spectrum from red (uncertain) to blue (predictable).

## Acknowledgments

We thank Bingni Brunton, Venkatesh Gopal, and Agnese Seminara for helpful discussion and members of the Gire lab for comments on the manuscript. We would also like to thank Sarahi Carolina Ponton Junes and Ryan Van Ort for assistance with data collection. Funding provided by NIH/NIDCD grant R00 DC013305 (DHG), the FACE foundation and by the University of Washington.

## Author contributions

DHG and BJJ initiated the study and designed the experiments. DHG, BJJ and SO built the automated navigation arena and wrote the behavioral monitoring software. BJJ and GL performed experiments. DHG performed analysis of behavioral data with input from BJJ and GL. DHG and BJJ generated all figures reporting experimental data. DHG and BJJ wrote the text. All authors provided editing suggestions for the figures and text.

## Methods

### Subjects

The experiments in this study were performed on 12 male Long-Evans rats, purchased from Charles River Labs and housed individually. All animals were maintained on a 12-hour light-dark schedule (lights on at 7:00am) with *ad libitum* access to water. After a weeklong habituation to the animal housing facility, all animals were then sustained at 85% of their free-feeding body weight in order to maintain motivation. All tests were performed between 9:00am and 6:00pm. To limit distal visual cues, all tests were performed under dim red light (~660 nm). All experimental procedures were approved by the Institutional Animal Care and Use Committee at the University of Washington.

### Testing Apparatus

The foraging arena was a large, fully enclosed open-field measuring 2m in length, 1m in width, and 1m in height. The frame of the arena was constructed from T-slotted aluminum railings. The sides of the arena were constructed from 1.27cm thick clear acrylic, while the ceiling was 0.635cm in thickness. The floor was a sheet of 0.0635cm thick opaque white acrylic. The ends of the arena were made from a wire mesh to allow for air to circulate throughout. A nest area where the animals would remain during the intertrial interval was attached to one end of the arena. The nest area was constructed from 1.27cm thick clear acrylic. Two synchronized cameras (The Imaging Source; DMK 23UP1300; frame rate 120 per second) were used to track the movement of the animals. An automated, custom-made pellet dispenser was used to bait the arena with 45mg sucrose pellets (Bio-Serv). An Arduino Uno controlled the movement of the motors running the pellet dispenser, allowing movement in the x- and y-coordinate plane.

#### Estimation of odor cues

Odor cue dispersal in the arena was directly measured using an ethanol source and miniature ethanol sensors that were scanned in a grid across the arena. The maximal signal detected at each sensor location over 30 seconds was normalized and reported in Supp. Fig. 2. There was no flow imposed on the arena, which limited the dispersal of airborne odor cues.

### Behavioral Paradigm

Before testing, all animals were habituated to the animal facility for 1 week. Animals then spent 2 days habituating to the attached waiting cage for ~15 minutes at a time. In order to motivate animals to return to the waiting cage, sucrose pellets were placed in the cage every 2 minutes when a 1 second, 1000Hz tone was played. They were then granted access to the test arena and were given 2-3 days to habituate to it. Animals were considered to have reached criterion when they were able to make 3 transitions between the waiting cage and test arena within 30 minutes.

Animals were placed into the waiting cage at the beginning of each testing session. Rats completed 1 session a day of 3 trials each. Before each trial, the automated pellet dispenser baited the arena with sucrose pellets organized into 3 clusters of approximately 3 pellets each. During foraging periods the dispenser was automatically lifted out of the arena so that the animals could not interact with it. Procedures differed only through the testing phase, when animals were assigned to forage within environments of high, medium, or low food location predictability. Animals trained on the environment with high food location predictability (n=4) were overtrained on a single distribution of pellet locations that stayed consistent across trials and sessions. Animals foraging in the environment with low food location predictability (n=4) were trained on unpredictable pellet distributions that changed across trials. All other animals (n=4) were trained on a moderately predictable distribution of pellet locations that changed slightly over time. All rats were given a maximum of 30 minutes to eat all of the sucrose pellets during the session. The entire testing period lasted for 30-35 days with approximately 5 sessions a week.

### Data and Statistical Analysis

No explicit power analysis was conducted in order to determine sample sizes. However, the number of animals used is consistent with experiments in the current literature. All analyses were conducted using MATLAB (MathWorks). A custom LabView (National Instruments) program was used to collect the behavioral data. Significant differences between groups were assessed with the Mann-Whitney U test followed by p-value adjustment with False Discovery Rate when multiple comparisons were made.

Predictability of pellet distributions was quantified using an across trial minimum distance metric, which, for each pellet in a given distribution reports the minimum distance from that pellet to all pellets in the immediately previous distribution. Relative entropy (RE) is equivalent to Kullback-Leibler Divergence and was calculated as: *RE*(*P||Q*) = ∫ *P*(*j*)log(*P*(*j*)/*Q*(*j*)) for all points j in the current trial’s probability density function (P) and the probability density function calculated from all previous trials (Q). Prior to calculating the RE all distributions were convolved with a smoothing function, which was an averaging filter of width = 10 cm. RE is reported in bits.

For establishing optimal pellet acquisition sequences for each distribution we used a genetic algorithm (available at: https://www.mathworks.com/matlabcentral/fileexchange/21198-fixed-start-open-traveling-salesman-problem-genetic-algorithm). Efficiency of foraging paths (Fig. 2e) was calculated as *fe = lo/la*, where *lo* is the optimal path length, *la* is the animal’s path length, and *fe* is foraging efficiency.

### Bayesian search

For analyses conducted in Figure 3 and Supplemental Figure 2, we modeled rat behavior as a Bayesian search. Briefly, the search arena is divided into 2.8 cm squares resulting in a 40 x 80 grid of possible locations. This grid is then populated with the same pellet distributions that were used in the behavioral experiments. We start our analysis on day 10 of training, which provides an agent with up to the first 10 days of training data as a map of prior expectations regarding pellet locations (see Supp. Fig. 2). The expression for prior expectation of pellet location is given by:

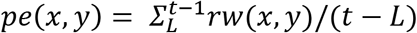

Where *t* is the trial number, *rw* is the probability of a pellet being found at a given point, (*x,y*), over previous trials and *pe* is the resulting prior expectation from the previous pellet locations. *L* is based on the length of memory being used and is defined as *L* = *max*(*t* — *md*, 1), with *md* being memory depth in trials, with *md* >=1. To enforce the nearest-neighbor search strategy used by rats, this map of prior expectations is discounted by linear distance from the agent, resulting in decreased likelihood to search first in areas that are located at large distances from the agent. This results in the following expression at a point, (*x, y*) within the grid of possible pellet locations:

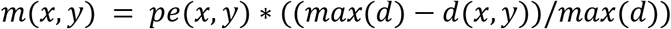

Where *d* is the distance from the agent and *m* is the memory-based map of prior expectations for pellet location adjusted by distance from the agent. The agent also uses sensory information that decays with distance to update their expectation of the possible pellet location,

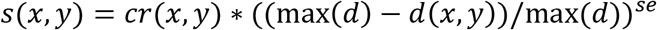

where *s* is the sensory density function and *cr* is a map with the current location of all pellets set to 1 and all other locations set to 0. The term *se* is an exponent that determines the rate of decay of sensory information with distance. These two sources of information are weighted and then summed to result in a map that guides the agent’s next step in the search path.

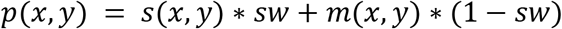

Where *p* is the probability map, *s* is the sensory density function and *m* is the memory-based map of prior expectations for pellet location. The term *sw* is the weight given to sensory information, {*sw* | 0 ≤ *sw* ≤ 1}. The agent makes its next step along the vector to the maximum point of *p*. The agent is considered to have perfect target detection at their location, such that after the agent moves to a new location, if a pellet is at that location it is always detected and if no pellet is at that location the probability of a target at that site is updated to 0. To fit parameters for the Bayesian search, we used a 3-dimensional coarse grid of values for *sw, se*, and *md*. We found the best fit for each animal in this grid and report these results in Fig. 3 and Supp. Fig. 2.

For reported measures in Figure 3c, *sa* = (1 – (*se/max*(*SE*)) + *sw*)/2, where *sa* is sensory acuity and *SE* is the set of values of *se* across all best fits for 12 animals, while 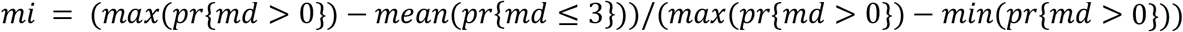, where *mi* is long-term memory usage and *pr* is the correlation of the agent’s performance with the animal’s performance using *md* set to the indicated range of values.

**Supplementary Fig. 1:**
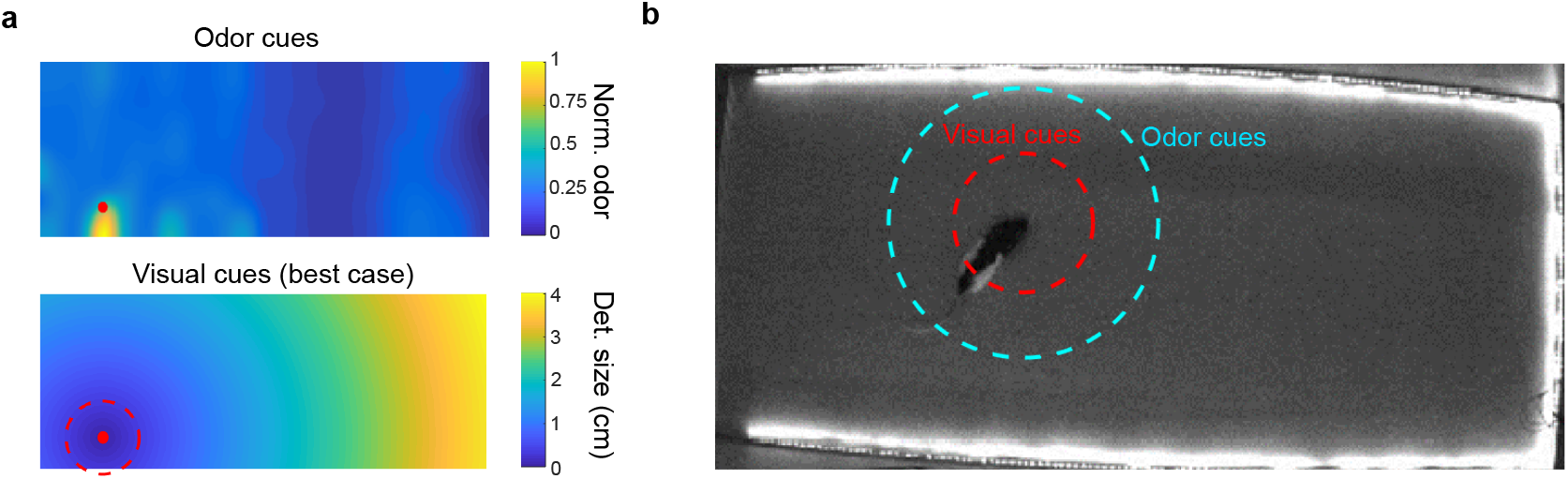
Sensory cues are local. **a) Top**: Experimentally determined spread of odor in the foraging arena (see methods). **Bottom**: Calculated size of a pellet necessary for it to be visible for a foraging rat under bright, broad-spectrum lighting conditions with high contrast, based on reported values for rat visual acuity^1,2^. The dashed red line indicates the actual size of the pellets used (and thus the distance for detection under ideal conditions). All experiments in the current study were done under dim red light using pellets matched in color to the arena floor, further limiting the range for visual detection. **b)** Estimated best-case pellet detection distances for olfactory (cyan) and visual (red) sensory cues. Due to both the dim, red lighting conditions and the lightly odorized pellets actual detection distances are likely to be much smaller.

**Supplementary Fig. 2:**
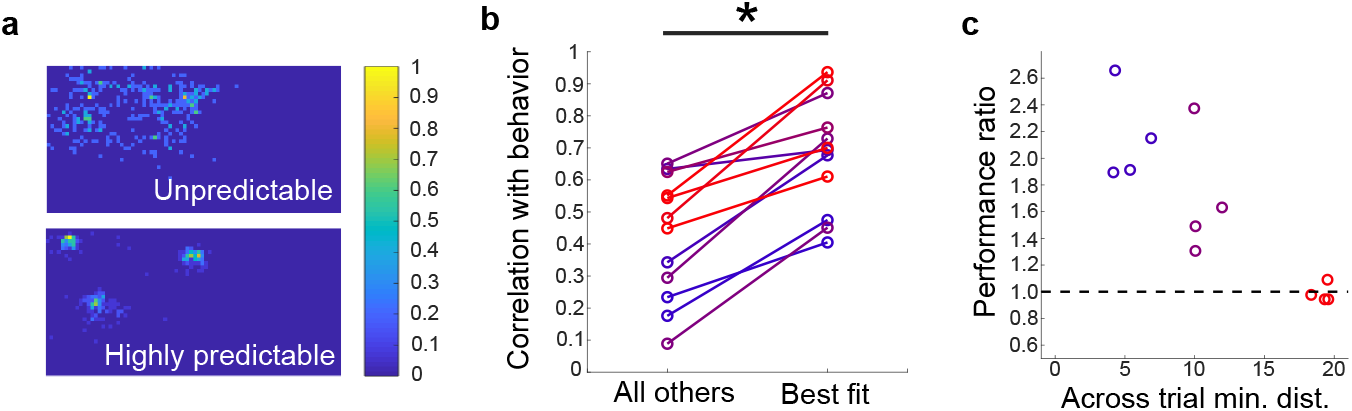
Modeling behavior as a Bayesian search. **a)** Examples of prior distributions accumulated over all trials for one predictable and one unpredictable set of pellet locations. **b)** Correlation of agent’s search performance with animal behavior when using parameters fit to other animals (All others) or the best fit to that specific animal (Best). The best fit is significantly better than the fits from other animals (p= 0.0043, n = 12). **c)** Performance ratio (Path length with priors from different distributions / Path length with correct prior) for all animals plotted as a function of the across trial minimum distance for the distributions presented to each animal (significant correlation: R = −0.85, p = 0.0004). A higher value for the performance ratio indicates longer path length with a shuffled prior. Agents searching with unpredictable distributions (red) show identical performance regardless of the prior used.

